# Macroscale traveling waves evoked by single-pulse stimulation of the human brain

**DOI:** 10.1101/2023.03.27.534002

**Authors:** Justin M. Campbell, Tyler S. Davis, Daria Nesterovich Anderson, Amir Arain, Zac Davis, Cory S. Inman, Elliot H. Smith, John D. Rolston

**Affiliations:** Interdepartmental Program in Neuroscience, University of Utah, Salt Lake City, UT, USA; Department of Neurosurgery, University of Utah School of Medicine, Salt Lake City, UT, USA; School of Biomedical Engineering, Faculty of Engineering, University of Sydney, Sydney, New South Wales, Australia; Department of Neurology, University of Utah, Salt Lake City School of Medicine, UT, USA; Department of Ophthalmology & Visual Sciences, University of Utah School of Medicine, Salt Lake City, UT, USA; Department of Psychology, University of Utah, Salt Lake City, UT, USA; Department of Biomedical Engineering, University of Utah, Salt Lake City, UT, USA; Department of Neurosurgery, Brigham & Women’s Hospital, Harvard Medical School, Boston, MA, USA

**Keywords:** Traveling wave, CCEP, SPES, stimulation

## Abstract

Understanding the spatiotemporal dynamics of neural signal propagation is fundamental to unraveling the complexities of brain function. Emerging evidence suggests that cortico-cortical evoked potentials (CCEPs) resulting from single-pulse electrical stimulation may be used to characterize the patterns of information flow between and within brain networks. At present, the basic spatiotemporal dynamics of CCEP propagation cortically and subcortically are incompletely understood. We hypothesized that single-pulse electrical stimulation evokes neural traveling waves detectable in the three-dimensional space sampled by intracranial stereoelectroencephalography. Across a cohort of 21 adult patients with intractable epilepsy, we delivered 17,631 stimulation pulses and recorded CCEP responses in 1,019 electrode contacts. The distance between each pair of electrode contacts was approximated using three different metrics (Euclidean distance, path length, and geodesic distance), representing direct, tractographic, and transcortical propagation, respectively. For each robust CCEP, we extracted amplitude-, spectral-, and phase-based features to identify traveling waves emanating from the site of stimulation. Many evoked responses to stimulation appear to propagate as traveling waves (∼14-28%), despite sparse sampling throughout the brain. These stimulation-evoked traveling waves exhibited biologically plausible propagation velocities (range 0.1-9.6 m/s). Our results reveal that direct electrical stimulation elicits neural activity with variable spatiotemporal dynamics, including the initiation of neural traveling waves.

**Significance Statement:** Using single-pulse stimulation, we identify a subset of intracranial evoked potentials that propagate as neural traveling waves. Our results were robust across a range of distinct but complementary analysis methods. The identification of stimulation-evoked traveling waves may help to better characterize the pathways traversed by spontaneous, pathological, or task-evoked traveling waves and distinguish biologically plausible propagation from volume-conducted signals.

## Introduction

With the advent of multi-sensor brain imaging, brain signals that had been assumed to be stationary have been revealed to propagate in space and time (Prechtl et al., 1997). These so-called neural traveling waves have gained increasing interest over the past decade (Muller et al., 2018; Foster and Scheinost, 2024). Neural traveling waves are ubiquitous, showing important roles during development (Schoch et al., 2018), sleep (Muller et al., 2016), epileptic activity (Alarcon et al., 1997; Smith et al., 2016, 2022; Martinet et al., 2017; Yearley et al., 2024), cognitive states (Zhang et al., 2018; Mohan et al., 2024), and evoked sensory activity (Muller et al., 2014; Zhaoyang et al., 2020; Aggarwal et al., 2022). Mesoscopic waves typically propagate at speeds from 0.1 to 0.8 meters per second, reflecting axonal conduction delays (Girard et al., 2001). More recently, waves have been shown to change with behavior and to modulate local neuronal firing (Davis et al., 2020). While traveling waves have generally been characterized in oscillations across a two-dimensional grid of sensors, they have also been observed in evoked potentials from sensory stimuli and in three-dimensional space (Aggarwal et al., 2022; Yearley et al., 2024).

Cortico-cortical evoked potentials (CCEPs) provide causal insight into the effective connectivity of the brain (Keller et al., 2014b). These responses to single pulses of electrical current applied directly to the human brain have been used for many different applications—for example, localizing epileptogenic tissue and seizure spread (Valentín et al., 2002; Lega et al., 2015; Matsumoto et al., 2017), mapping distinct types of connectivity (Entz et al., 2014; Keller et al., 2014a; Crocker et al., 2021a), quantifying signal complexity across levels of consciousness (Comolatti et al., 2019; Arena et al., 2021, 2022; Mofakham et al., 2021), and tracking the effects of stimulation on neural plasticity (Keller et al., 2018; Huang et al., 2019). Researchers have begun to investigate the temporal properties of CCEPs, characterizing the relationship between the amplitude and latency of the early N1 component (10-50 ms) and axonal conduction delays resulting from the underlying white matter architecture (Trebaul et al., 2018; Silverstein et al., 2020; Lemaréchal et al., 2021). For example, a recent study that combined CCEPs with tractography reported considerable age-related changes in the transmission speed of cortico-cortical signals (Blooijs et al., 2023). Emerging evidence suggests that the temporal dynamics of CCEPs may also provide novel insights into the patterns of information flow between and within brain networks (Veit et al., 2021; Seguin et al., 2023).

In this study, we employed contemporary methods to test the hypothesis that single pulse stimulation of the human brain evokes neural traveling waves. We found that, indeed, a large proportion of CCEPs met the definition of traveling waves and characterized their patterns of propagation across the brain surface and through white matter tracts. Our findings were robust across distinct but corroborative analysis methods that capture different characteristics of the evoked response (i.e., amplitude, spectral power, phase). Taken together, these results highlight a previously uncharacterized response to direct electrical stimulation. Showing that CCEPs behave as traveling waves opens the possibility of using these evoked potentials to characterize the pathways trod by spontaneous, pathological, or task-related traveling waves. Further, biologically plausible propagation may be used as a method of discriminating authentic, local CCEPs from artifactual or volume-conducted CCEPs, improving the spatial and temporal localization of these electrophysiological markers.

## Materials and Methods

### Participants

Twenty-one adult patients with medically intractable epilepsy (n = 7 female) underwent intracranial monitoring with stereoelectroencephalography (SEEG) to localize their seizure foci (**Table 1**). We performed stimulation mapping while patients were off their antiseizure medications during their inpatient hospital stay. Results from some of these stimulation sessions were reported in prior studies of evoked potential recordings (Kundu et al., 2020; Davis et al., 2021a). All patients provided informed consent prior to participation, and our team took care to implement the recommended principles and practices for ethical intracranial neuroscientific research (Feinsinger et al., 2022). The study was approved by the University of Utah Institutional Review Board.

**Table 1.** Demographics and clinical characteristics of the patient cohort. SPES = single-pulse electrical stimulation, SOZ = seizure onset zone, OFC = orbitofrontal cortex, MFG = middle frontal gyrus, PVNH = periventricular nodular heterotopia.

| **Patient ID** | **Age (yrs.)** | **Sex** | **Electrode (no.)** | **Stimulated (no., %)** | **SPES (no.)** | **SOZ** |
| --- | --- | --- | --- | --- | --- | --- |
| P1 | 25.5 | M | 56 | 56 (100%) | 448 | Bilateral mesial temporal lobe |
| P2 | 25.9 | M | 97 | 94 (96.9%) | 940 | None identified |
| P3 | 35.3 | F | 81 | 81 (100%) | 648 | Right hippocampus |
| P4 | 44.3 | M | 69 | 69 (100%) | 966 | Right hippocampus |
| P5 | 42.4 | F | 48 | 48 (100%) | 480 | Right OFC |
| P6 | 42.2 | F | 54 | 54 (100%) | 540 | None identified |
| P7 | 35.6 | F | 28 | 24 (85.7%) | 240 | Right MFG |
| P8 | 45.0 | M | 52 | 52 (100%) | 520 | None identified |
| P9 | 41.7 | F | 30 | 23 (76.7%) | 1150 | Bilateral mesial temporal lobe |
| P10 | 29.1 | F | 39 | 28 (71.8%) | 1400 | Bilateral mesial temporal lobe |
| P11 | 33.2 | M | 40 | 36 (90.0%) | 1800 | Left temporal |
| P12 | 27.5 | M | 46 | 38 (82.6%) | 1634 | Right OFC |
| P13 | 22.7 | M | 48 | 44 (91.7%) | 880 | Right mesial temporal |
| P14 | 54.0 | M | 20 | 20 (100%) | 600 | Right PVNH |
| P15 | 30.3 | M | 40 | 32 (80.0%) | 960 | Left mesial temporal |
| P16 | 29.8 | M | 49 | 43 (87.8%) | 1161 | Left frontal operculum |
| P17 | 59.6 | M | 36 | 31 (86.1%) | 930 | Left temporal |
| P18 | 43.2 | M | 34 | 34 (100%) | 374 | Bilateral hippocampi |
| P19 | 45.1 | F | 42 | 39 (92.9%) | 780 | Left hippocampus |
| P20 | 26.5 | M | 55 | 38 (69.1%) | 760 | Bitemporal |
| P21 | 37.7 | M | 51 | 30 (58.8%) | 330 | Bilateral hippocampi |

### Electrode Localization and Registration

To localize electrodes, we co-registered each patient’s preoperative magnetization-prepared rapid gradient-echo (MP-RAGE), fluid-attenuated inversion recovery (FLAIR), or T1 MRI to their post-operative CT using custom MATLAB software and the *Statistical Parametric Mapping Toolbox* (SPM 12; https://www.fil.ion.ucl.ac.uk/spm/software/spm12/). Automated image processing, electrode detection, and anatomical localization of intracranial electrodes were performed using the open-source *Locate electrodes Graphical User Interface* (LeGUI; https://github.com/Rolston-Lab/LeGUI) software (Davis et al., 2021a) and the Brainnetome atlas (Fan et al., 2016). Electrode contact locations are visualized in MNI space using the *Visbrain* software package (Combrisson et al., 2019) (**Fig. 1A**).

**Fig 1.**
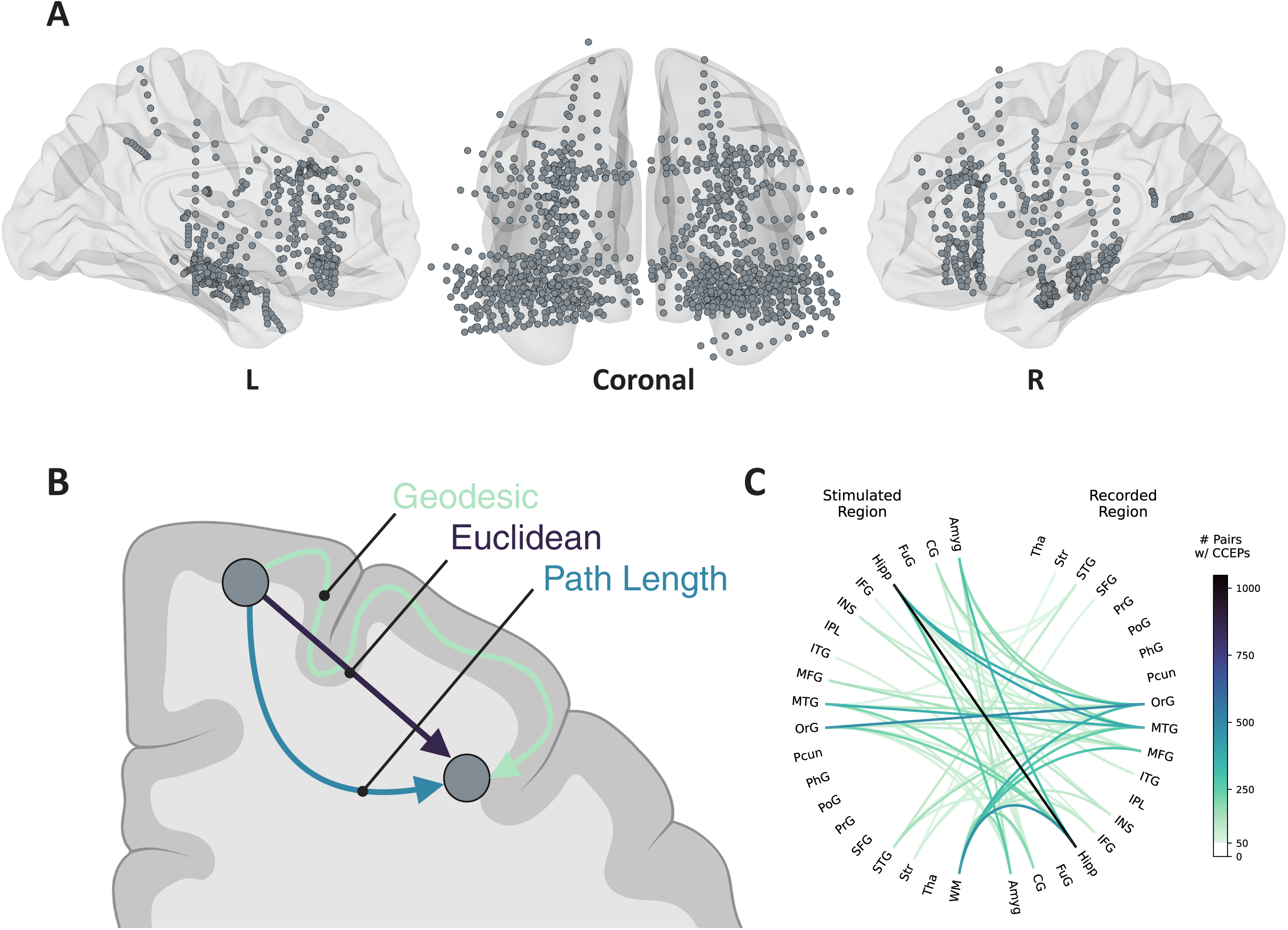
Electrode contact locations, distance metrics, and stimulation summary. **A.** Brain models depicting SEEG electrode contact locations in MNI space with midsagittal and coronal views (n = 21 patients). **B.** Schematic comparison of distance measurements between two hypothetical electrode locations. Euclidean distance (dark purple) represents the shortest straight-line distance between two points. Path length (blue) represents the shortest distance between two points along each patient’s individualized MRI tractography map. Geodesic distance (green) represents the shortest distance between two points along a path restricted within the cortex. **C.** Chord diagram illustrating the regions where stimulation was applied and where corresponding CCEP responses were recorded. Lines are colored by the number of unique stim-response contact pairs with robust CCEPs (lines with < 50 contact pairs are not shown). Hipp = hippocampus, WM = white matter, MTG = middle temporal gyrus, OrG = orbitofrontal gyrus, Amyg = amygdala, MFG = middle frontal gyrus, CG = cingulate gyrus, INS = insula, Str = striatum, STG = superior temporal gyrus, IFG = inferior frontal gyrus, ITG = inferior temporal gyrus, PrG = precentral gyrus, FuG = fusiform gyrus, SFG = superior frontal gyrus, IPL = inferior parietal lobule, PoG = postcentral gyrus, PhG = parahippocampal gyrus, Pcun = precuneus, Tha = thalamus.

### Electrophysiological Recordings and Stimulation

Neurophysiological data were recorded from intracranial SEEG electrodes with variable contact spacing (Ad-Tech Corp., Racine, WI; DIXI Medical, France) using a 128-channel neural signal processor (Neuroport, Blackrock Microsystems, Salt Lake City, UT) and sampled at 1kHz. Following recommended practices, we chose an intracranial electrode in the white matter with minimal artifact and epileptiform activity as the online reference (Mercier et al., 2022). Stimulation was delivered using a 96-channel neurostimulator (Cerestim, Blackrock Microsystems, Salt Lake City, UT).

CCEP mapping consisted of monopolar single-pulse electrical stimulation (biphasic, 5.0-7.5 mA, 500 ms pulse width) administered to a subset of electrodes in a randomly selected order every 2.5-3.5 s over an ∼45 min session (Kundu et al., 2020). Each electrode contact received between 4-50 discrete trials of single pulse electrical stimulation; variability in trial count across contacts stemmed from time constraints and clinical considerations (i.e., emphasis on hypothesized seizure foci).

### Signal Processing

Intracranial recordings were first divided into pre-stimulation and post-stimulation epochs (-1000 to -5 ms and 5-1000 ms, respectively), with stimulation onset set at time zero. Peri-stimulation data (-5 to 5 ms) was removed to mitigate stimulation artifacts. Next, data were bandpass filtered at 0.3-250 Hz and re-referenced offline to the common median across channels to minimize noise (Rolston et al., 2009).

We subsequently measured the broadband, high-gamma power (70-150 Hz) envelope from local field potentials recorded during the post-stimulation epochs to characterize the spectrotemporal response to single pulses of electrical stimulation. Specifically, we bandpass filtered the post-stimulation data using a zero-phase fourth-order Butterworth filter, applied the Hilbert transform, and multiplied the result by its complex conjugate.

### Distance Metrics

We used three distinct measures of distance to capture subcortical and cortical traveling waves in 3-dimensional (3D) SEEG space: Euclidean distance, path length, and geodesic distance (**Fig. 1B**). These metrics were chosen to compare biologically plausible metrics (i.e., path length, geodesic distance) against naïve approaches which do not account for the brain’s unique geometry (i.e., Euclidean distance).

#### Euclidean

The Euclidean distance between each electrode pair is defined by the length of the shortest line segment connecting the two electrode locations in 3D space. Accordingly, we calculated the pairwise 3D Euclidean distance between all possible contacts using the following:

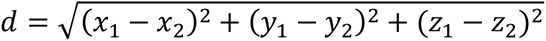

where *x_n_*, *y_n_*, and *z_n_* correspond to the 3D anatomical coordinates of the electrode contact within the patient’s brain.

#### Path length

The path length is the average distance between two electrodes derived from patient-specific tractography. Diffusion-weighted imaging (DWI) was acquired for all patients as part of standard clinical care. Using the *FMRIB* Software Library (FSL), DWI images were eddy corrected, and a network connectivity matrix was generated using network mode in the *probtractx2* function (Behrens et al., 2007; Charlebois et al., 2022). A volume with a 0.5 cm radius at each electrode centroid was used as the seed region, and segmentation was computed in *FreeSurfer* (Fischl, 2011; Anderson et al., 2021). Finally, the output N x N network matrix, which corresponds to average tractographic distances between each combination of N number of contacts, was made symmetric.

#### Geodesic

The geodesic distance is the shortest distance along the cortical surface between two electrodes, following gyri and sulci. Cortical surface segmentations for each hemisphere were generated within *FreeSurfer* using each patient’s structural MRI (Fischl, 2011). Using these surfaces, the geodesic distance was calculated with the *SurfDist* python package (v 0.15.5, https://pypi.org/project/surfdist/) between each electrode pair on the same hemisphere. Electrodes were excluded from this distance calculation if that electrode’s centroid was more than 1 cm from the nearest cortical surface (33.7% of stim-response contact pairs).

### CCEP Detection

Fundamental considerations in the detection and analysis of CCEPs have been reviewed previously (Prime et al., 2018). Numerous methods for CCEP detection have been employed in prior studies, including amplitude thresholding (Crocker et al., 2021b), machine-learning frameworks analyzing evoked response shapes (Miller et al., 2021, 2023), stimulation-induced gamma-based network identification (SIGNI) (Crowther et al., 2019), and others (Prime et al., 2020b).

In this study, evoked responses were considered CCEPs if (1) the trial median of the early response period (5-100 ms post-stimulation) contained ≥ 15 ms of continuous values more than threefold greater than the median (across time and trial) of the immediate pre-stimulation epoch (-100 to -5 ms) and (2) the median (across time and trial) of the early response period (5-100 ms post-stimulation) was > 30 µV, as previously described (Kundu et al., 2020). Responses measured from contacts located within white matter were excluded. Evoked responses that did not meet these criteria were excluded from subsequent analyses.

### CCEP Feature Extraction

To characterize patterns of macroscale CCEP propagation, we adapted methods used to analyze evoked traveling waves emanating from interictal epileptiform discharges (maximal descent) (Smith et al., 2022) and to characterize the phase of broadband oscillations in local field potentials (generalized phase) (Davis et al., 2020).

#### Maximal descent

For each CCEP identified, we determined the time at which the evoked response was most rapidly decreasing (i.e., maximal descent, MD), using:

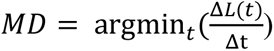

Where *L* represents the CCEP local field potential amplitude or high-gamma power (**Fig 2A-B**). Max descent was chosen, among other possible CCEP features (e.g., N1/N2 peak latency), because prior research has validated the feature as a computationally efficient and robust predictor for traveling wave analyses (Liou et al., 2017), and because CCEP morphology can be highly variable, often without canonical N1/N2 components (Miller et al., 2021, 2023).

**Fig 2.**
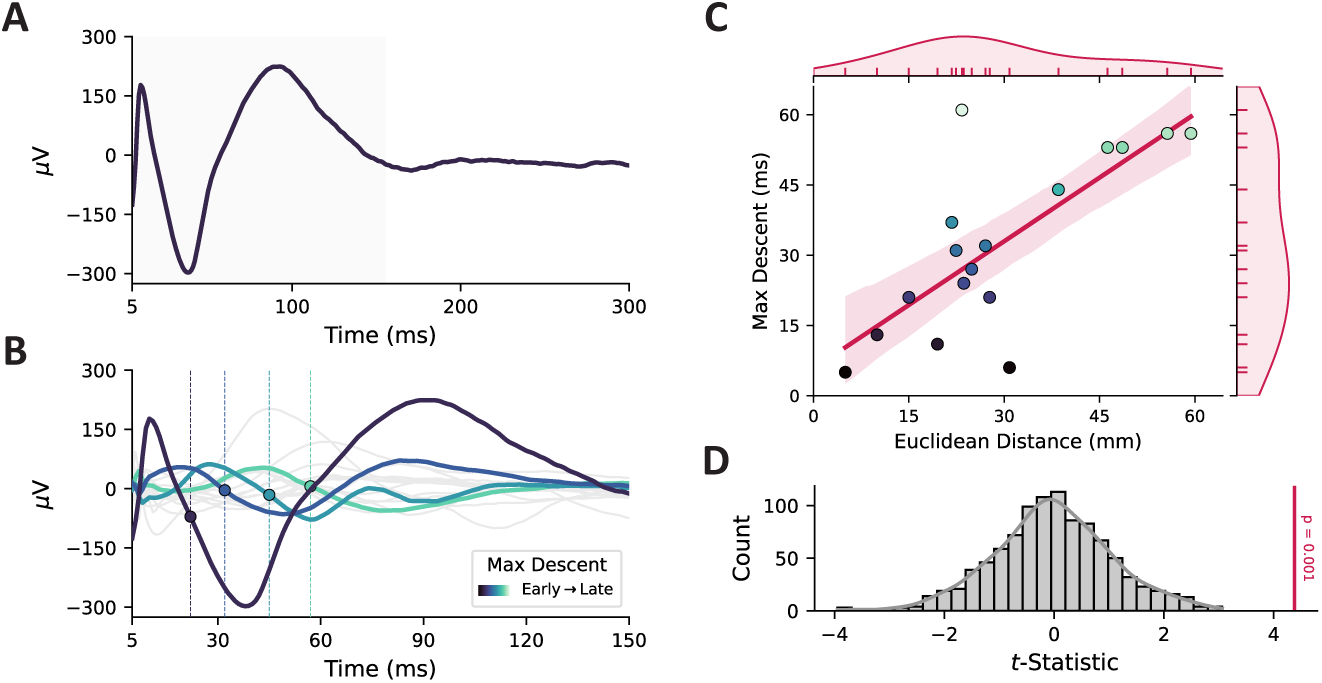
Traveling wave identification using max-descent features. **A.** Example CCEP evoked by single-pulse electrical stimulation. **B.** Subset of CCEPs with amplitude max descents (MD) marked. Stimulation at this location evoked robust CCEPs in several other electrodes, some of which are highlighted to show variability in max descent time (indicated with a circle and dashed line). **C.** Example of significant traveling wave. The distance (Euclidean shown) between electrodes was linearly regressed onto amplitude MD time to identify traveling waves. Dots represent electrodes with robust CCEPs, colored by time to MD (purple → green, early → late). The shaded area represents 95% CI. Kernel density estimate plots along the axes represent the distributions of distance and MD time. **D.** Example 1,000-fold permutation test comparing linear model *t*-values against a null distribution of shuffled distances.

#### Generalized phase

The generalized phase (GP) is a recently developed method that analyzes the phase of non-stationary broadband neural signals while overcoming the limitations of narrow-band filtering (Davis et al., 2020). Briefly, the evoked response is filtered in a wide bandpass to exclude low-frequency content (5-40 Hz), the Hilbert Transform is applied to the broadband signal, phase is computed using the four-quadrant arctangent function, and negative frequencies are corrected via cubic interpolation. The result captures the phase of the largest fluctuation on the recorded electrode. Finally, we extracted the average GP from the evoked response in a time window containing the empirical median CCEP amplitude max descent time (35-45 ms post-stimulation) (**Fig. 3A-B**).

**Fig 3.**
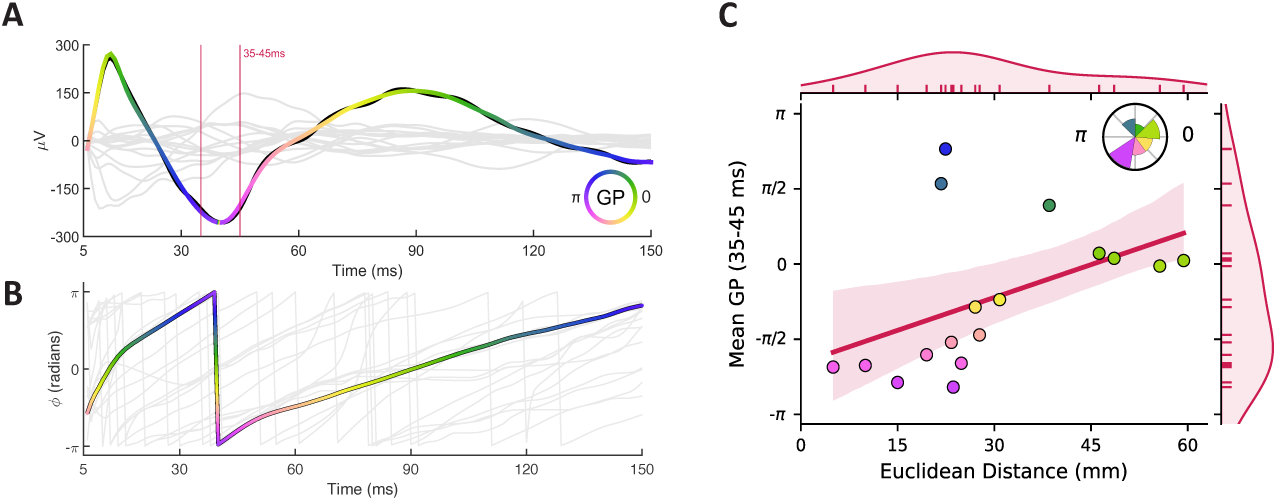
Traveling wave identification using generalized phase. **A.** Example CCEPs evoked by electrical stimulation. The trial-averaged response from a single contact is highlighted; the line color represents the generalized phase (GP) value at each point in time. Red lines highlight the time window used to extract the average GP for regression (10-20 ms). **B.** Corresponding phase values for each of the recorded CCEPs, spanning -π to +π. **C.** Linear regression of Euclidean distance onto GP reveals a robust traveling wave (p < 0.001). The circular histogram (panel inset, top right) shows the distribution of GP values from CCEPs included in the model.

### Traveling Wave Identification and Characterization

For each contact stimulated, we regressed the distance (Euclidean, path length, or geodesic) between each stimulation contact and the contacts where robust CCEPs were recorded onto their respective analysis metric (amplitude MD time, high-gamma MD time, GP). A stimulation contact was determined to evoke a traveling wave if a significant linear relationship between distance and analysis metric was observed across response contacts; to control for false positives and identify robust model fits, we performed 1000-fold permutation testing against a null distribution of shuffled distances (**Fig. 2C-D**, **Fig. 3C**). Only the linear models with a test statistic at the p < 0.05 level compared to the permuted distribution were considered traveling waves. The propagation velocity of traveling waves identified from amplitude or high-gamma MD time was obtained from the slope of the linear model and converted to m/s to facilitate comparison with what has been reported previously. To estimate propagation velocity in the phase-based models, we calculated the average instantaneous frequency during the 35-45 ms post-stimulation period, took the median instantaneous frequency across response contacts, multiplied the result by 2π, and divided by the slope of the regression model.

### Experimental Design and Statistical Analyses

We used a linear mixed-effects model to analyze the effect of stimulated region, distance (Euclidean, EUC; path length, PL; geodesic, GEO), and analysis metric (amplitude MD time, AMP; high-gamma MD time, HG, generalized phase, GP) on the proportion of stimulated contacts which evoked traveling waves; patient ID was included as a random effect due to the variability in traveling waves elicited across patients (**Supplemental Fig. 1**). Pairwise post-hoc testing of distance and analysis metrics was performed using Tukey’s or Dunn’s tests. Given the large number of regions stimulated, we chose to further characterize the proportions of traveling waves expected by stimulating inside vs. outside each region using a series of chi-squared tests for independence followed by false discovery rate correction.

To characterize differences in model fit and traveling wave velocity, we first performed Anderson-Darling tests for normality (Anderson and Darling, 1952), which determined that non-parametric statistical tests were most appropriate. Accordingly, we used a two-way ANOVA on ranks to compare median model R^2^ values (distance metric and analysis metric as factors) and pairwise Kolmogorov-Smirnov tests to compare the distributions of traveling wave velocities across distance metrics. Statistical analyses were performed using the *scipy* (Virtanen et al., 2020) and *statsmodels* (Seabold and Perktold, 2010) libraries. Data were visualized using *matplotlib* (Hunter, 2007)*, seaborn* (Waskom, 2021), and *MNE* (Gramfort et al., 2014).

## Results

Across all patients, we recorded from 1,019 intracranial electrode contacts (48.5 ± 17.5 per patient, mean ± SD) (**Fig. 1A**). We administered a total of 17,631 trials of single pulse electrical stimulation (839.6 ± 421.7 per patient) to 917 intracranial electrode contacts (43.7 ± 19.0 per patient) (**Fig. 1C**). The demographics and clinical characteristics of the patient cohort are shown in **Table 1**.

We observed robust traveling waves across all analysis metrics (amplitude MD time, AMP; high-gamma MD time, HG, generalized phase, GP) and distance metrics (Euclidean, EUC; path length, PL; geodesic, GEO) (**Fig. 4A**; example shown in **Supplemental Video 1**). The linear mixed effects model revealed significant differences in the proportion of traveling waves evoked across region stimulated and distance metrics while accounting for individual differences (p < 0.05). There were no significant differences in the proportion of traveling waves identified across analysis metrics. Pairwise post-hoc testing revealed that the mean proportion of traveling waves measured with EUC was greater than PL (mean difference = 6.4%, 95% CI [2.8%, 9.9%], p = 0.001), PL was more common than GEO (mean difference = 5.0%, 95% CI [1.5%, 8.5%], p = 0.003), and EUC was more common than GEO (mean difference = 11.3%, 95% CI [7.8%, 14.9%], p < 0.001).

**Fig 4.**
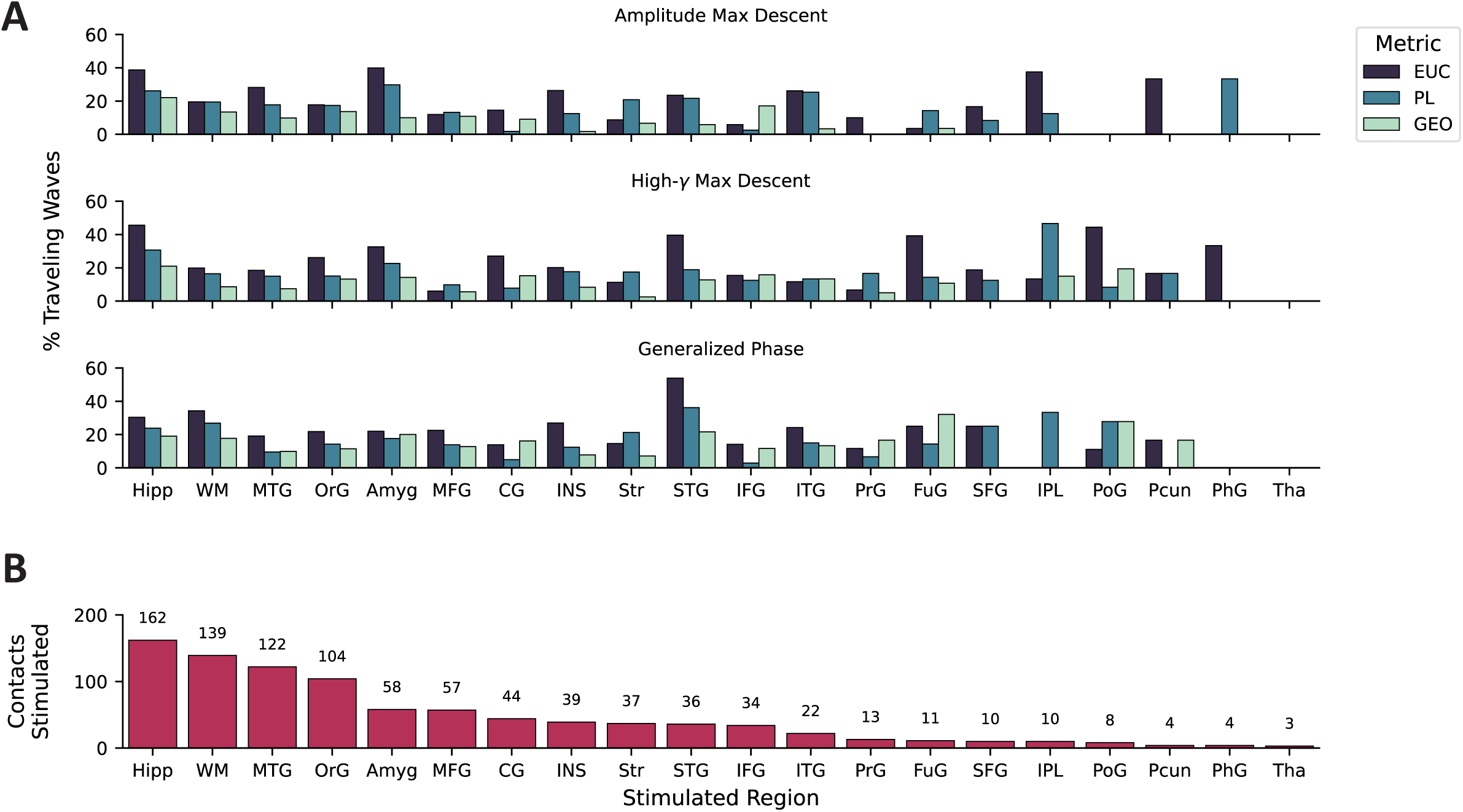
Proportion of traveling waves evoked by stimulating different regions. **A.** Bars represent the proportion of stimulated contacts which evoked a traveling wave (%), separated by region (n = 20), distance metric, and analysis metric. **B.** The number of unique contacts stimulated in each region (ordered high → low). EUC = Euclidean (purple), PL = path length (blue), GEO = geodesic (green).

Since stimulation was applied to areas distributed throughout the brain, we next sought to characterize regional differences in the proportion of CCEPs that appeared as evoked traveling waves. The electrode localization and registration pipeline identified stimulated electrodes located in 20 distinct regions (**Fig. 4B**). Chi-square tests comparing the proportion of traveling waves resulting from stimulation inside vs. outside the region of interest showed a greater proportion of traveling waves evoked from stimulation in the hippocampus (Hipp), superior temporal gyrus (STG), and white matter (WM) relative to other brain regions with EUC (Hipp: χ^2^(1, 917) = 46.0, p < 0.001; STG: χ^2^(1, 917) = 8.0, p = 0.029; WM: χ^2^(1, 917) = 6.6, p = 0.044), for the hippocampus, white matter, and amygdala (Amyg) with PL (Hipp: χ^2^(1, 917) = 43.8, p < 0.001; WM: χ^2^(1, 917) = 7.8, p = 0.029; Amyg: χ^2^(1, 917) = 6.4, p = 0.044), and for the hippocampus and amygdala with GEO (Hipp: χ^2^(1, 917) = 23.5, p < 0.001; Amyg: χ^2^(1, 917) = 7.8, p = 0.029). A smaller proportion of traveling waves than would be expected was observed in the middle frontal gyrus (MFG), superior temporal gyrus (STG), striatum (Str), and inferior frontal gyrus (IFG) with EUC (MFG: χ^2^(1, 917) = 17.0, p = 0.001; Str: χ^2^(1, 917) = 10.9, p = 0.01; IFG: χ^2^(1, 917) = 7.1, p = 0.039), the cingulate gyrus (CG), middle inferior gyrus, inferior frontal gyrus, and middle temporal gyrus with PL (CG: χ^2^(1, 917) = 13.2, p = 0.003; MFG: χ^2^(1, 917) = 6.4, p = 0.044; IFG: χ^2^(1, 917) = 9.8, p = 0.015; MTG: χ^2^(1, 917) = 6.1, p = 0.047), and the striatum and middle frontal gyrus with GEO (χ^2^(1, 917) = 8.0, p = 0.029; MFG: χ^2^(1, 917) = 6.6, p = 0.044).

By measuring the slope of the regression, we were able to estimate the propagation velocity of each traveling wave identified; to minimize the effect of velocities that were biologically implausible, we excluded values that were negative or were outside the range defined by 3x the median absolute deviation (Leys et al., 2013). A series of Kolmogorov-Smirnov tests comparing the velocity distributions observed across distance metrics revealed that EUC < GEO < PL for AMP (EUC vs. GEO: D = 0.6, p < 0.001; EUC vs. PL: D = 0.8, p < 0.001; PL vs. GEO: 0.5, p < 0.001) HG (EUC vs. GEO: D = 0.6, p < 0.001; EUC vs. PL: D = 0.8, p < 0.001; PL vs. GEO: 0.3, p < 0.001), and GP (EUC vs. GEO: D = 0.4, p < 0.001; EUC vs. PL: D = 0.7, p < 0.001; PL vs. GEO: 0.5, p < 0.001) (**Fig. 5**). The median (Q1-Q3) propagation velocities of traveling waves identified with AMP were 1.0 (0.8-1.3) m/s, 2.0 (1.3-2.6) m/s, and 3.1 (2.4-4.0) m/s for EUC, GEO, and PL, respectively; we observed similar, albeit slightly slower, velocities with HG: (EUC: 0.8 (0.5-1.1) m/s, GEO: 1.5 (1.1-2.1) m/s, PL: 2.2 (1.5-3.2) m/s). In contrast, propagation velocities estimated from GP models were faster than those calculated for AMP and HG, across all distance metrics (EUC: 1.5 (1.0-2.1) m/s, GEO: 2.4 (1.8-3.3) m/s, PL: 4.5 (2.7-5.9) m/s).

**Fig 5.**
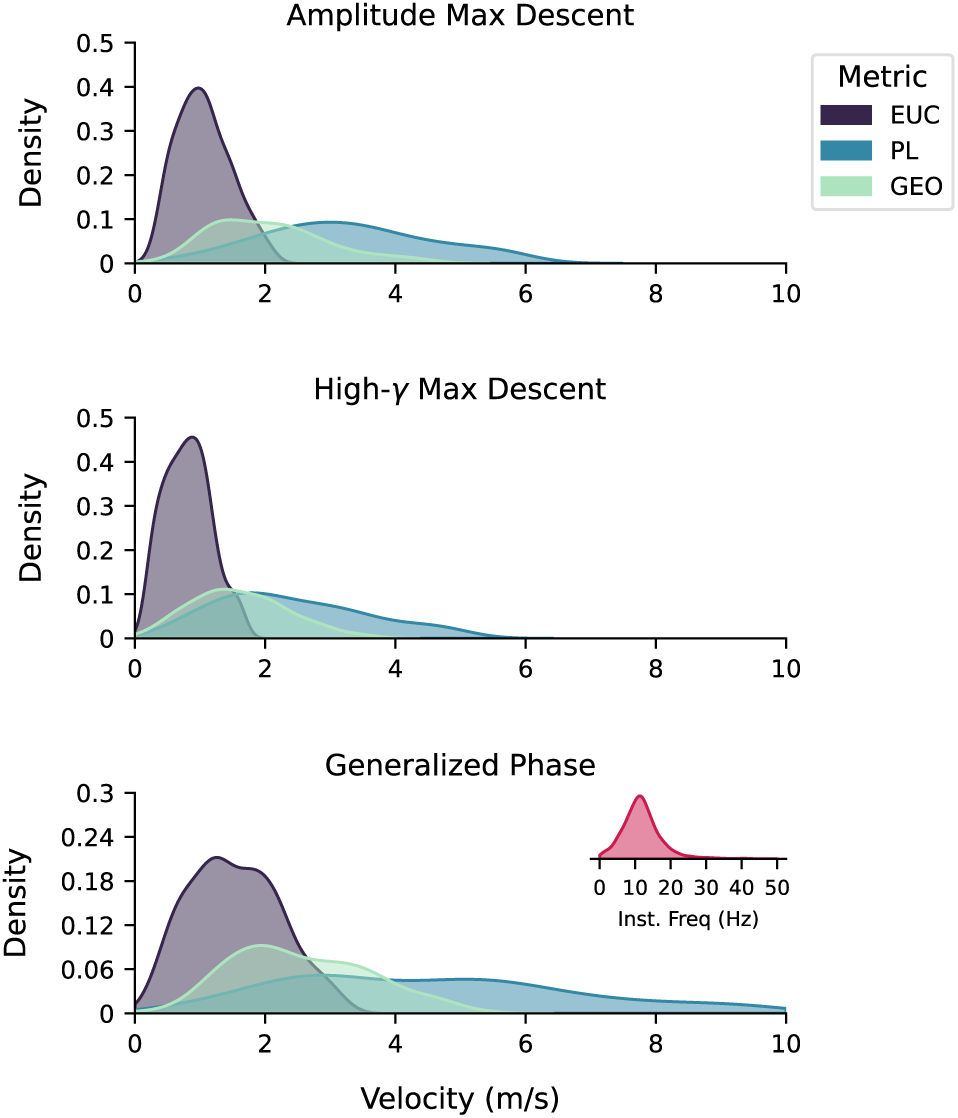
Gaussian kernel-density estimate of the traveling wave velocities observed using the analysis metrics (amplitude max descent, high-gamma max descent, generalized phase); colors represent distance metrics. EUC = Euclidean (purple), PL = path length (blue), GEO = geodesic (green). The inset on the bottom panel shows the distribution of instantaneous frequencies observed across all responses during the 35-45 ms generalized phase analysis window.

The median time to max descent in the CCEPs was 38.0 ms (± 34.6) and 25.0 ms (± 44.1) for evoked amplitude and high-gamma power, respectively. Max descent timings across the two measures of amplitude and high-gamma power were positively correlated [r(13583) = 0.26 (p < 0.001)] (**Fig. 6A**). To further characterize model performance, we compared the proportion of variance explained (R^2^) using a two-way ANOVA on ranks, with distance and analysis metrics as factors. We observed a significant main effect of distance metric (F(2, 1574) = 25.6, p < 0.001), but not for analysis metric (F(2, 1574) = 2.9, p = 0.054). A post-hoc Dunn’s test revealed that models using GEO had a median R^2^ value greater than that of EUC (p < 0.001) and PL (p = 0.002), and that EUC > PL (p = 0.001) (**Fig. 6B, Supplemental Fig. 2**). Exploratory analyses comparing the proportion of stimulation contacts mutually identified across analysis metrics suggests that AMP, HG, and GP are largely capturing distinct features of the evoked response (**Fig. 6C**).

**Fig 6.**
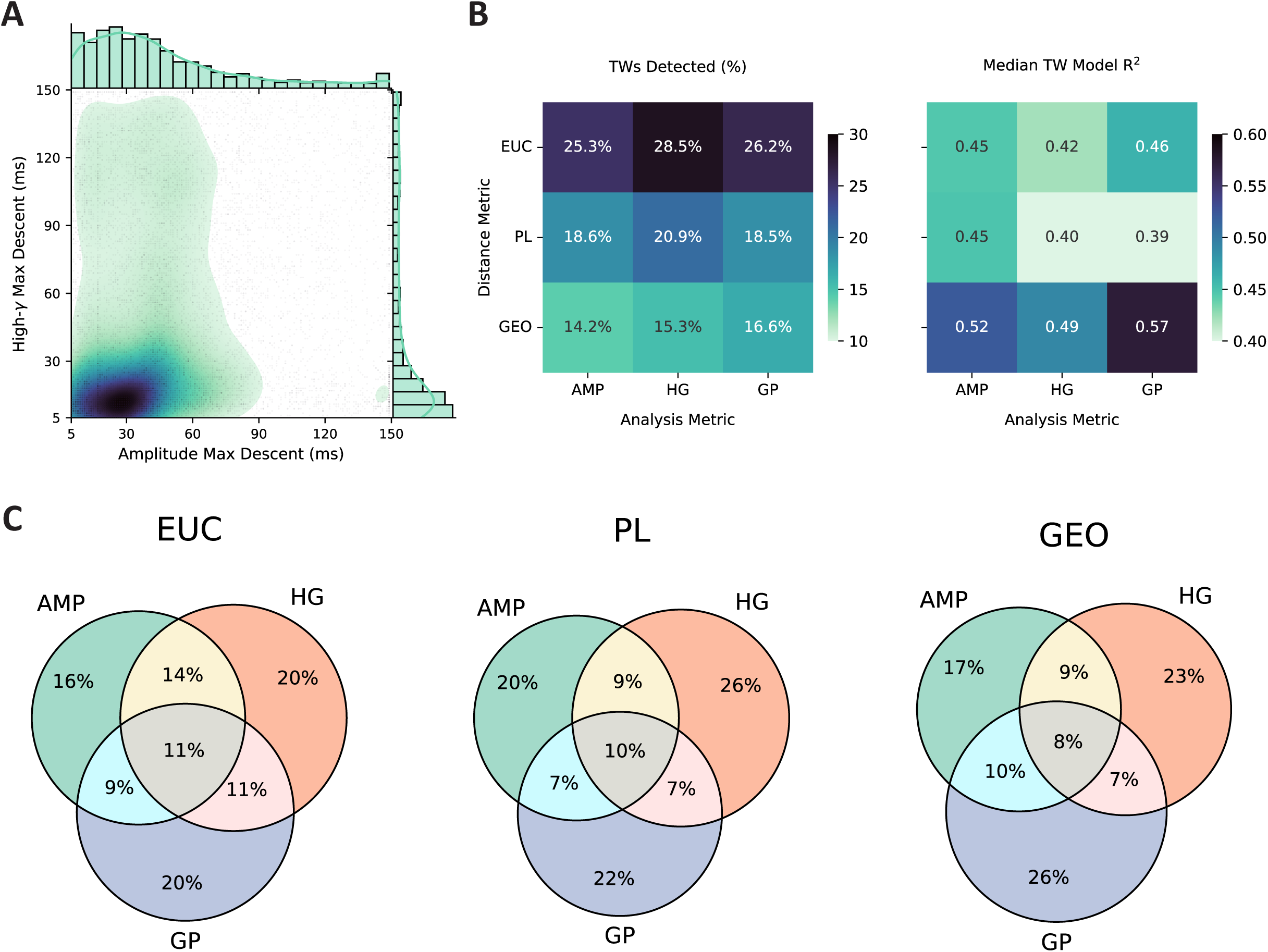
Comparison of different metrics used for traveling wave identification. **A.** Scatterplot of evoked amplitude and high-gamma (70-150 Hz) MD times with overlaid Gaussian kernel-density estimate. The shaded area represents the probability of values at each location (green → dark purple, low → high; probabilities of < 20 % are not shown). Histograms along axes show the distributions of MD times. **B.** Heatmaps of the proportion of traveling waves detected (left) and model R^2^ values across distance and analysis metrics. **C.** Venn diagrams showing the proportion of traveling waves mutually identified across different distance and analysis metrics. EUC = Euclidean, PL = path length, GEO = geodesic, AMP = amplitude MD time, HG = high-gamma MD time, GP = generalized phase.

## Discussion

We applied single-pulse electrical stimulation to intracranial SEEG electrodes in a cohort of patients with intractable epilepsy to test the hypothesis that direct electrical stimulation evokes neural traveling waves. In doing so, we observed a subset of CCEPs that propagate as neuronal traveling waves and exhibit similar characteristics to traveling waves observed in spontaneous and task-evoked activity.

The range of propagation velocities we observed (0.1-9.6 m/s) was comparable to those reported previously for mesoscopic (0.1-0.8 m/s) and macroscopic traveling waves (1-10 m/s)– consistent with the axonal conduction velocities of unmyelinated and myelinated white matter fibers within the cortex, respectively (Muller et al., 2018). Large-scale network models suggest that neural traveling waves emerge spontaneously from these axonal conduction delays (Davis et al., 2021b). Similar traveling wave velocities have also been reported in the human brain from microelectrode array recordings of IEDs (Smith et al., 2022) and spontaneous neocortical alpha and theta oscillations (Zhang et al., 2018). Additionally, a recent study that leveraged biologically informed modeling of CCEPs and tractography from patients in the F-TRACT database (f-tract.eu) reported a median cortico-cortical axonal conduction velocity of 3.9 m/s (Lemaréchal et al., 2021), a value very close to what we observed in our analogous AMP-PL models of traveling wave propagation along white matter tracts: 3.1 m/s.

We recorded traveling waves in ∼14-28% of the channels stimulated, depending on the detection metrics used. There are several reasons why stimulation-evoked traveling waves may not have been observed more ubiquitously. For one, we utilized SEEG recordings, which inherently provide a sparse representation of 3D volumes and the cortical surface, since contacts along the same lead are arranged linearly (Anderson et al., 2021). Moreover, SEEG contact orientation relative to the nearest neural source may add an additional source of variance, compared to consistently oriented surface electrodes. Subsequent investigation of stimulation-evoked traveling wave dynamics in high-density electrocorticographic recordings would offer superior spatial resolution for estimating wave speed and shape. Additionally, since the shape of CCEP responses can be highly variable, metrics like max descent may fail to generalize to less stereotypical response shapes, necessitating the use of phase-based metrics like GP.

Methodological constraints aside, the observation that only a subset of CCEPs exhibit traveling wave characteristics raises an interesting question: Are spatiotemporal delays in CCEP propagation a possible means of separating biologically plausible evoked responses from spurious signals? Volume conduction, for example, is a commonly observed artifact in evoked responses to stimulation that confounds the interpretation of CCEPs (Prime et al., 2020a). With our approach, volume-conducted signals could be distinguished from genuine traveling waves because they lack a robust, linear relationship between CCEP max descent and electrode distance. Although not the focus of this study, further characterization of CCEPs as a multiregional response may provide novel, clinically relevant insights for the mapping of epileptogenic networks (Bernabei et al., 2023).

Robust CCEPs were most often observed near the stimulation site, adjacent to the stimulating electrode, or along the same lead. These closely spaced electrodes may be best characterized by Euclidean distance, which could explain why this metric led to a higher proportion of traveling waves identified relative to path length or geodesic distance, metrics which are more biologically plausible paths for subcortical and cortical traveling wave propagation. That notwithstanding, models using geodesic distance appear to explain a greater proportion of variance than those using Euclidean distance or tractographic path length. This suggests that stimulation-evoked traveling waves are best modeled via propagation along the cortical surface.

Our comparison of the regional differences in the response to stimulation suggests that stimulation-evoked traveling waves are more common from particular brain regions, like the hippocampus and amygdala; previous studies have reported that single pulse stimulation of these areas tends to elicit robust, broadly distributed CCEPs (Enatsu et al., 2015; Mégevand et al., 2017; Guo et al., 2021). Prior studies have observed traveling waves in areas throughout the human neocortex (Zhang et al., 2018). Given that medial temporal lobe structures tend to be oversampled relative to other brain regions, the increased electrode density may have some relationship to the regional differences in stimulation-evoked traveling waves that we observed.

The study has a few noteworthy limitations. First, electrode locations are determined solely by clinical considerations, leading to heterogeneity between patients and a denser sampling of regions known to be highly epileptogenic (e.g., mesial temporal lobe). Our prior work on traveling waves was performed using Utah electrode arrays, which have the advantage of a grid-like orientation, allowing a more straightforward characterization of the spatiotemporal dynamics of traveling wave propagation.

Although max descent is a validated measure for traveling wave analyses (Liou et al., 2017), it reduces the entirety of the CCEP to a single, discrete measurement with respect to time. Other statistical approaches that leverage the high sampling rate of continuous time-series recordings (e.g., phase-based analyses) have been used in prior studies of endogenous traveling waves (Halgren et al., 2019; Davis et al., 2020). For this reason, we additionally characterized the generalized phase of the evoked response. With both approaches, we observed statistically significant stimulation-evoked traveling waves. These results suggest that this approach to traveling wave detection is robust and that our observations are not simply an artifact of data processing.

Finally, we restricted the CCEP feature extraction window to 5-150 ms post-stimulation for our analyses. This window was chosen to include the first downward deflection typically observed in our population of CCEPs. In other studies, this has been referred to as N1 (10-50 ms) or the early negative deflection of the slower N2 (50-200 ms) component. When these separate components are observable, recent evidence suggests they may represent intra- and internetwork communication, respectively (Veit et al., 2021). Other analyses have shown that morphological characteristics of evoked waveforms, outside the traditional N1/N2 paradigm, may offer more powerful insights into the anatomy underlying the observed CCEPs (Miller et al., 2021).

Subsequent studies of stimulation-evoked traveling waves may benefit from analyzing the evoked responses from these two components separately or leveraging the full response profile of the evoked signal.

## Conclusions

Our analyses of CCEPs revealed that many evoked responses to single-pulse electrical stimulation propagate as traveling waves. The stimulation-evoked traveling waves we observed were robust across multiple distinct methods of identification and exhibited characteristics in line with what has been reported previously for spontaneous traveling waves. Traveling waves were observed across distributed brain areas but were most frequently evoked when stimulating highly connected hub regions (e.g., hippocampus). These results highlight previously uncharacterized spatiotemporal dynamics of signal propagation in response to stimulation.

## Acknowledgments

Figure 1B was created with BioRender

## Abbreviations

CCEP: cortico-cortical evoked potential
EUC: Euclidean distance
iEEG: intracranial electroencephalography
GEO: geodesic distance
GP: generalized phase
MD: maximal descent
PL: path length
SEEG: stereoelectroencephalography
SPES: single-pulse electrical stimulation

## Disclosures

JDR has received prior consulting fees from Corlieve Therapeutics, Medtronic, NeuroPace, and Turing Medical.

## Funding

JMC was supported by the National Institute of Neurological Disorders and Stroke (T32 NS115723). JDR was supported by an NIH/NINDS career development award (K23NS114178). DNA was supported by an NIH/NINDS award (F32NS114322). CSI was supported by an NIH/NIMH R01 (R01MH120194) and NSF Foundations Grant (NSF2124252).

**Supplemental Fig 1.** Proportion of traveling waves evoked by stimulation in each patient, collapsed across regions. EUC = Euclidean, PL = path length, GEO = geodesic.

**Supplemental Fig 2.** Gaussian kernel-density estimate of the linear model R^2^ values across distance and analysis metrics. EUC = Euclidean, PL = path length, GEO = geodesic.

**Supplemental Video 1.** Illustrative example of a stimulation-evoked traveling wave vs. null response to single-pulse stimulation in a single patient. Patient-specific 3D brain models are shown with superimposed electrode contact locations. Stimulation of contact 5 (caudate) evoked a traveling wave (left), whereas stimulation of contact 16 (middle temporal gyrus) resulted in a null response indicative of volume conduction (right); stimulated contacts are shown in red. The max descent (MD) time of each robust CCEP is highlighted at the moment it occurs (yellow). Lower panels show robust CCEPs resulting from stimulation.

## Notes

Conflicts of Interest: **None**

### Summary of Updates

Substantive edits throughout. The introduction has been rewritten to contextualize prior investigations of traveling waves. New phase-based analysis methods have been added for traveling wave identification and characterization.

