## Supplementary figures and images for "Macroscale traveling waves evoked by single-pulse stimulation of the human brain"

### Supplemental Fig 1

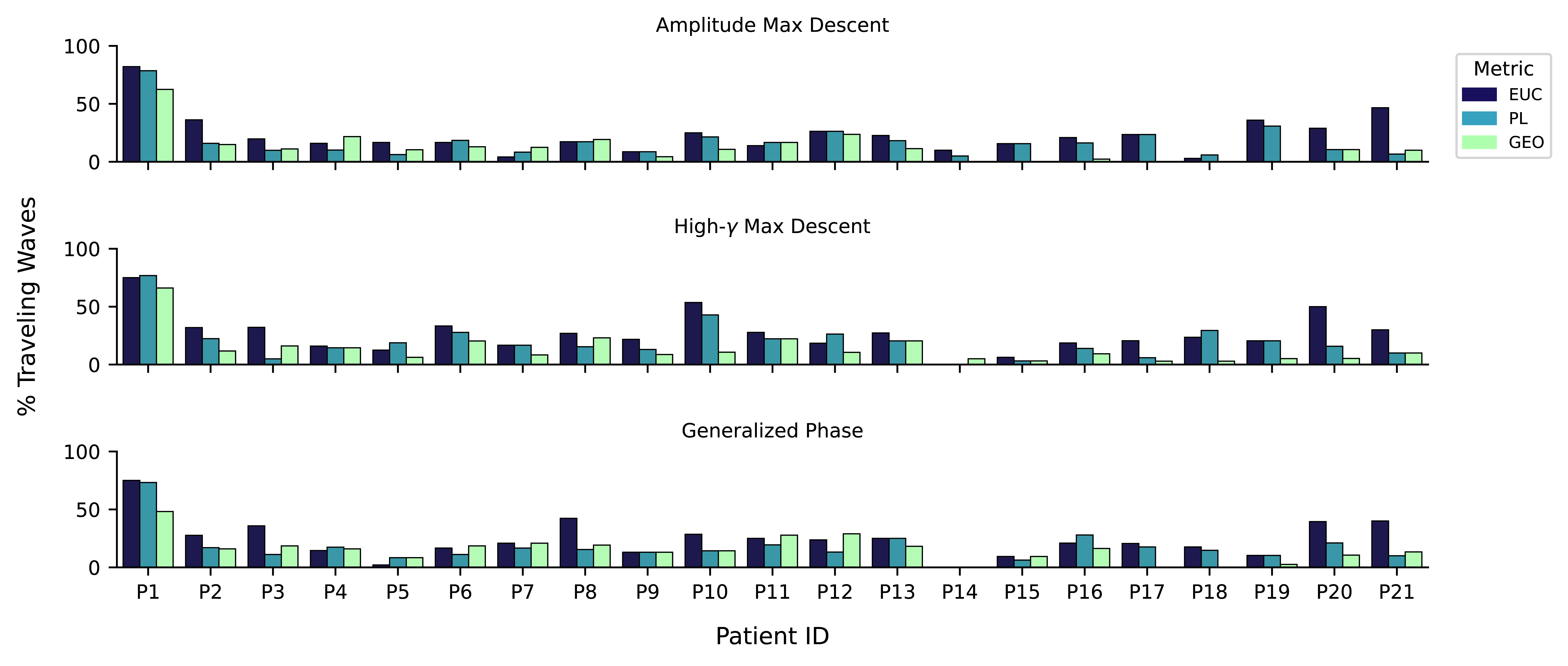

### Supplemental Fig 2

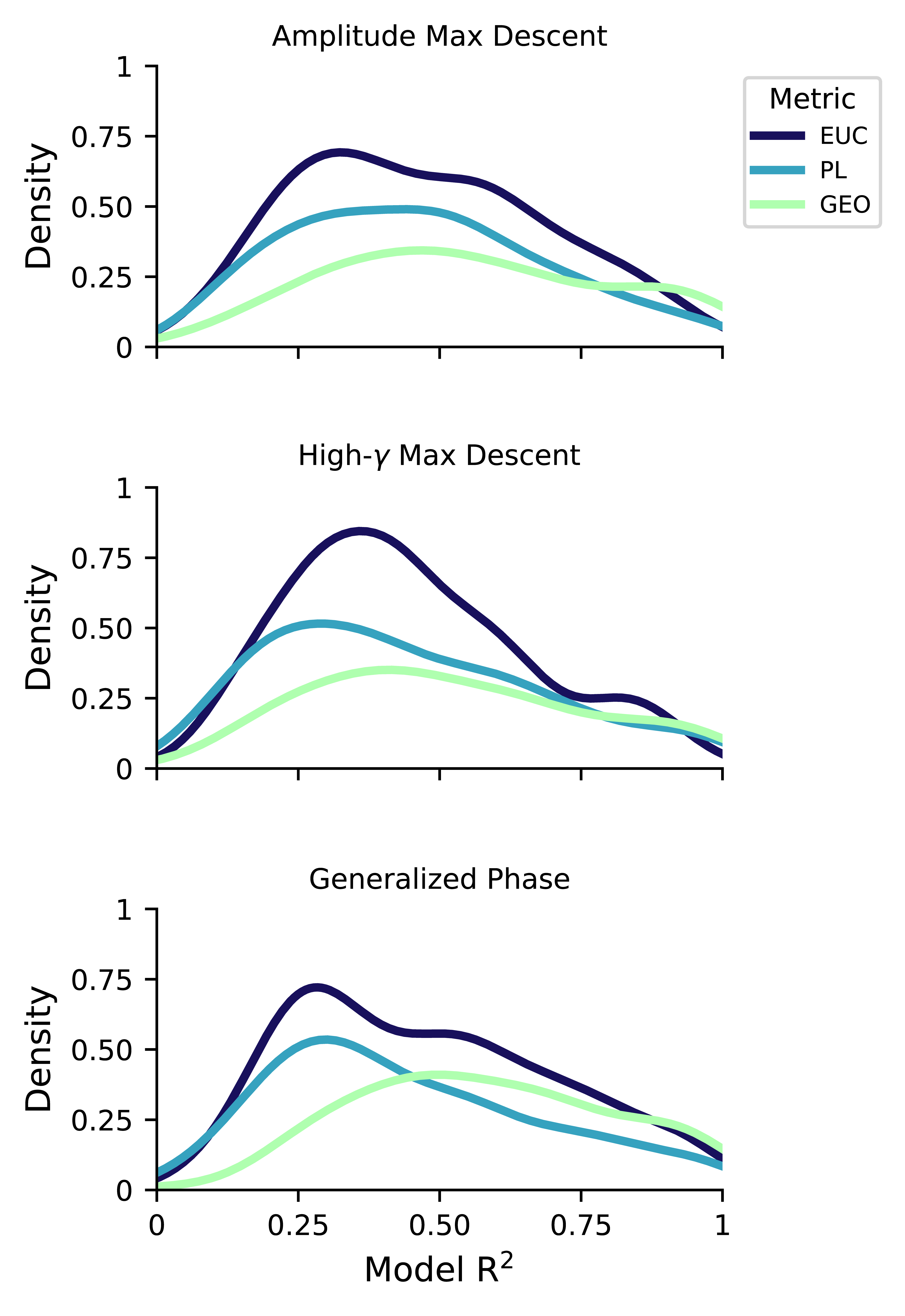
